# Optogenetic translocation to subcellular compartments through regulation of protein avidity

**DOI:** 10.1101/2025.04.23.650348

**Authors:** Zikang (Dennis) Huang, Yueying Gu, Yuzhi (Carol) Gao, Alexander Byrd, Hana Bader, Lukasz J. Bugaj

**Affiliations:** Department of Bioengineering, University of Pennsylvania, Philadelphia, PA, 19104, USA; Institute for Regenerative Medicine, University of Pennsylvania, Philadelphia, PA, 19104, USA; Abramson Cancer Center, University of Pennsylvania, Philadelphia, PA, 19104, USA

## Abstract

Inducible translocation to subcellular compartments is a common strategy for protein switches that control a variety of cell behaviors. However, existing switches achieve translocation through induced dimerization, requiring constitutive anchoring of one component into the target compartment and optimization of relative expression levels between the two components. We present a simpler, single-component strategy called Avidity-assisted targeting (Aviatar). Aviatar achieves translocation with only a single protein by converting low-affinity monomers into high-avidity assemblies through inducible clustering. We demonstrated the Aviatar concept and its generality using optogenetic clustering to drive translocation to the plasma membrane, endosomes, golgi, endoplasmic reticulum, and microtubules using binding domains for lipids or endogenous proteins that were specific to those compartments. Aviatar recruitment regulated actin polymerization at the cell periphery and revealed compartment-specific signaling of receptor tyrosine kinase fusions associated with cancer. Finally, GFP-targeting Aviatar probes allowed inducible localization to any GFP-tagged target, including endogenously-tagged stress granule proteins. Aviatar is a straightforward platform that can be rapidly adapted to a broad array of targets without the need for their prior modification or disruption.

## Introduction

Recruitment of proteins to specific subcellular compartments is a common mode of cellular regulation, for example in mitogenic signaling or in cytoskeletal assembly^1–3^. Many probes for on-demand control of cell behavior regulate translocation in response to external inputs including chemicals, light, or temperature^3–7^. These systems commonly use inducible dimerization: a cytoplasmic protein is recruited to a binding partner that is constitutively anchored in the target compartment. However, dimerization-based systems require multiple components that may need optimization of relative levels, which can be tedious or impractical depending on the experimental system^8^. Furthermore, a permanent exogenous anchor could disrupt the native function of the target compartment.

Single-component translocation probes would address these concerns, though few exist. Such probes currently either require sequence-specific insertion of a light-sensing domain into a binding domain^9^, or they require the binding domain to be masked and unmasked with light, which works best for only small binding peptides^10^. An alternative strategy can be found in the naturally-evolved BcLOV4 photoreceptor from *Botrytis cinerea*^11^. Upon light stimulation, BcLOV4 clusters and subsequently translocates to the plasma membrane ^11–14^. Clustering drives BcLOV4 translocation by increasing avidity of an otherwise low-affinity membrane-binding fragment, a concept observed across many natural proteins^15^. In principle, such regulated avidity could be broadly applied for probes that translocate to arbitrary cell compartments in a straightforward and modular manner.

Here, we present such a strategy called <u>Avi</u>dity-<u>a</u>ssisted <u>tar</u>geting (Aviatar). Aviatar is a suite of single-component probes that translocate to subcellular compartments including the plasma membrane (PM), golgi, endoplasmic reticulum (ER), endosome, microtubules (MT), and stress granules (SG), all through regulated avidity of domains that bind endogenous lipids or proteins. Aviatar probes allowed light-induced, compartment-specific regulation of the actin cytoskeleton and receptor tyrosine kinases, and activation across different compartments recapitulated known effects of protein mislocalization in human cancer. Finally, a GFP-targeting Aviatar allowed localization to any protein tagged with GFP, providing a flexible way to target arbitrary compartments even when binding domains cannot be easily found.

## Results

### Avidity can regulate translocation of proteins to the plasma membrane

Aviatar probes are composed of two modular functional domains: 1) a binding domain with weak affinity for a target compartment (wBD), and 2) a protein that can be clustered with an external input like light (e.g. Cry2 or its variants, **Figure 1A**)^16–19^. Upon light stimulation, clustering generates a high-avidity interaction, causing translocation of the clustered assembly. To demonstrate this concept, we first used amphipathic helix 1(AH1) from BcLOV4 as a wBD. AH1 mediates membrane-binding of BcLOV4 but by itself is too weak to drive localization^11,14^. Accordingly, the AH1-mCh-Cry2 fusion was diffuse in its dark, monomeric form. However, light stimulation induced rapid translocation to the membrane (**Figure 1B-E**), whereas a control construct lacking AH1 clustered in the cytoplasm. Notably, membrane translocation of AH1-mCh-Cry2 was found at all expression levels tested, including at exceedingly low levels where Cry2 clusters would normally be too small to be seen (**Figure S1**)^8^. Thus, AH1-Cry2 can implement a PM-targeting Aviatar probe (Aviatar_PM_). Aviatar_PM_ could also be generated with alternative wBD sequences, including the polybasic (PB) domain of STIM1 (**Figure 1F-G**), as shown previously^20^, as well as with the pleckstrin homology (PH) domain from Dynamin-1(shown using Cry2_olig_, **Figure 1H-I**)^21^.

**Figure 1.**
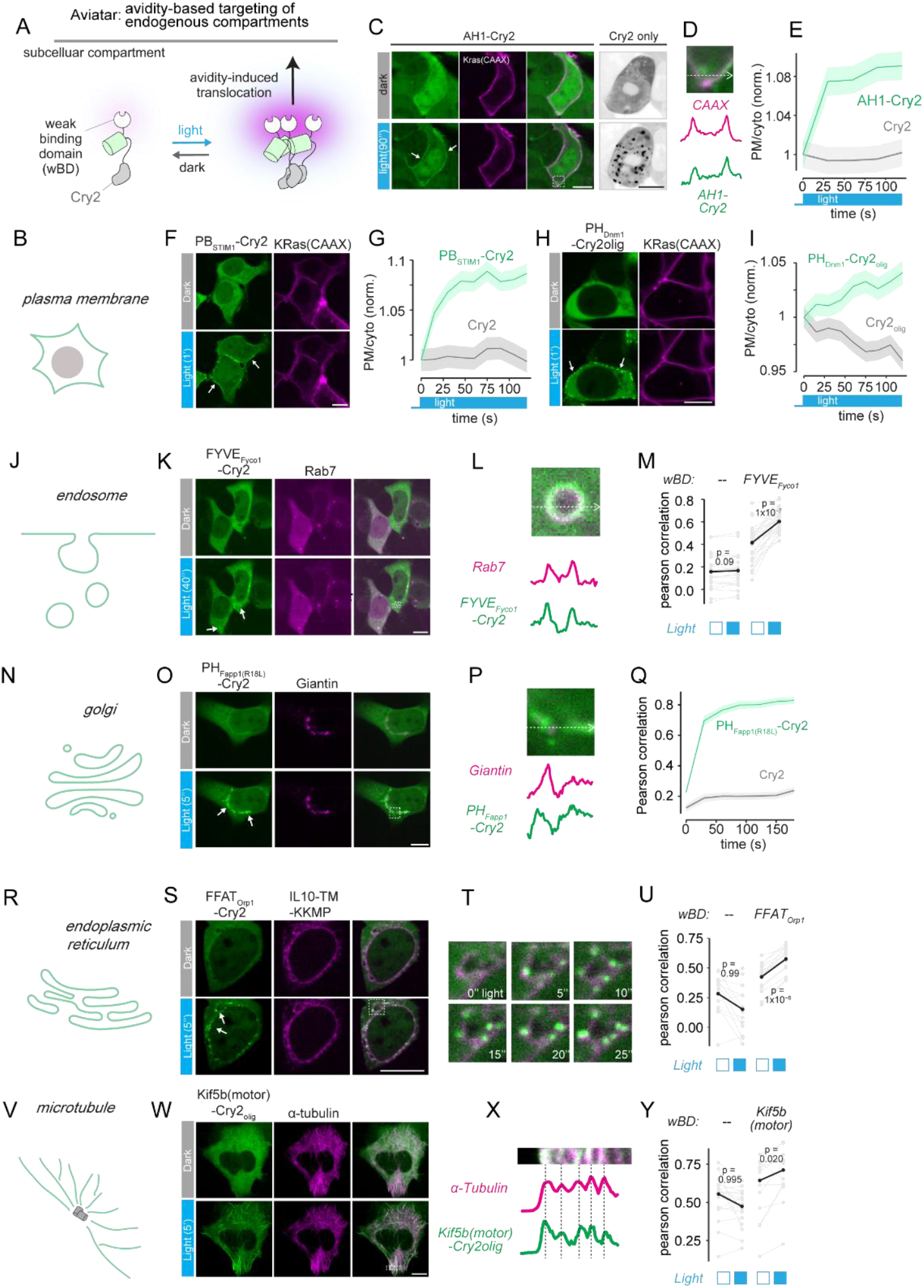
Aviatar can trigger translocation to arbitrary cellular compartments. **A**. Schematic of the Aviatar mechanism. A weak binding domain (wBD) is fused with Cry2. Upon light stimulation, increased avidity of wBD enhances its binding capability towards its targets. **B**. Aviatar controls translocation to the plasma membrane (PM). **C**. Upon light stimulation, AH1-Cry2 rapidly translocates to PM while Cry2 alone clusters in the cytoplasm. An iRFP-CAAX construct is used as a PM marker. Arrows highlight PM binding of Aviatar after light stimulation. **D**. Intensity profile of the highlighted region in (**C**). **E**. Quantification of (**C**). The ratio of mean intensity of PM and whole cell is normalized to the dark state (t = 0). **F**. Aviatar_PM_ demonstrated with PB_STIM1_ as the wBD. **G**. Quantification of (**F**), as described in (**E**). **H**. Aviatar_PM_ demonstrated with PH_Dnm1_ as the wBD, fused with Cry2olig. **I**. Quantification of (**H**), as described in (**E**). **J)** Aviatar controls translocation to endosomes. **K**. Aviatar_Endosome_ demonstrated using FYVE_Fyco_ as the wBD. Rab7 is used as an endosome marker. Arrows highlight localization of Aviatar to endosomes after light stimulation. **L**. Intensity profile of the endosome within the highlighted region of (**K**). **M**. Quantification of (**K**). Pearson correlation coefficient of the intensity of Aviatar and endosome marker channel in single cells. The same cells before and after light stimulation are connected by grey lines and the average is shown in black. **N**. Aviatar controls translocation to the golgi. **O**. Aviatar_Golgi_ demonstrated using PH_Fapp1_ as the wBD. Giantin is used as a golgi marker. Arrows highlight the localization of Aviatar to golgi after light stimulation. **P**. Intensity profile of the dashed region in (**O**). **Q**. Quantification of (**O**). Pearson correlation coefficient of the intensity of Aviatar_Golgi_ and golgi marker channel in single cells. **R**. Aviatar controls translocation to the ER. **S**. Aviatar_ER_ demonstrated using FFAT_Orp1_ as the wBD. A fusion protein composed of IL10 signal sequence, transmembrane domain, and ER retention signal KKMP is used as an ER marker. Arrows highlight the localization of Aviatar to ER after light stimulation. **T**. Enrichment of Aviatar on the ER over time in the highlighted region in (**S**). **U**. Quantification of (**S**). Pearson correlation coefficient of the intensity of Aviatar and ER marker channel in single cells is calculated. **V**. Aviatar controls translocation to microtubules. **W**. Aviatar_MT_ demonstrated using Kif5b(motor) as a wBD fused with Cry2olig, in HeLa cells. α-Tubulin is used as a microtubule marker. **X**. Intensity profile of the highlighted region in (**W**). **Y**. Quantification of (**W**). Pearson correlation coefficient of the intensity of Aviatar and the microtubule marker channel in single cells is calculated. Scale bars, 10 µm. All experiments were performed in HEK 293T cells unless otherwise specified. All line plots show mean +/- SEM of data.

### Generation of Aviatar toolkit for targeting arbitrary cell compartments

We next tested whether Aviatar could be generalized to target other subcellular compartments. Late endosomes and lysosomes are enriched for phosphatidylinositol-3-phosphate (PI(3)P)^22^, which is commonly leveraged for localization by proteins containing PI(3)P-binding FYVE domains^22–24^. We tested an Aviatar_endosome_ construct using the FYVE domain from Fyco1 (**Figure 1J-M**). Indeed, FYVE_Fyco1_-mCh-Cry2 appeared diffuse in the dark but showed clear localization to Rab7-labelled endosomes after light stimulation.

Aviatar_golgi_ probes were designed using a wBD for the golgi-specific protein ARF. We harnessed the PH domain from Fapp1, which, when harboring an R18L mutation, requires multimerization to bind ARF^25^. In HEK 293T cells, PH_FAPP1(R18L)_-mCh-Cry2 was cytoplasmic in the dark but colocalized with the golgi marker Giantin upon light stimulation (**Figure 1N-Q**). In a similar manner, we constructed an Aviatar_ER_ probe using an FFAT motif from the Orp1 protein, which binds to the ER-resident VAP (Vesicle-associated membrane protein (VAMP)-associated proteins) protein^26^. FFAT_Orp1_-mCh-Cry2 was diffuse in the dark but showed strong enrichment at the ER after light stimulation^27^(**Figure 1R-U**). Notably, the activated state appeared more granular than for other Aviatar probes, potentially due to the enrichment of VAP at membrane contact sites between organelles^28^.

Finally, we asked whether Aviatar could target non-membranous compartments such as cytoskeletal networks. Many proteins that associate with microtubules (MTs) are multimeric, though whether multimerization plays a causal role in microtubule binding is unclear^29–31^. We created successful Aviatar_MT_ probes by fusing microtubule-interaction regions from either Kif5B (1-325) or EML4(81-207), in the absence of neighboring multimerization sequences^29,32,33^. These constructs were both predominantly cytoplasmic in the dark but rapidly colocalized with fluorescently-labeled α-tubulin in the light (**Figure 1V-Y, S2**). These results validate Aviatar_MT_ and suggest a causal role for avidity in the normal MT-binding ability of the native proteins.

In summary, Aviatar is a general strategy that can be applied for single-component localization to various cellular compartments given the availability of a specific and weakly interacting binding domain.

### Aviatar probes can regulate cell physiology

To determine whether Aviatar probes can control cellular activity, we first tested regulation of events that had previously been controlled using dimerization-based recruitment. We asked if Aviatar_PM_ could regulate cytoskeletal networks near the plasma membrane through recruitment of the the DH-PH domain of Intersectin1 (DHPH_Itsn1_)^3,34–36^. Stimulation of Aviatar_PM_-DHPH_Itsn1_ induced rapid actin polymerization, indicated by filopodial and lamellipodial protrusions (**Figure 2A-C**). Protrusions were not observed in cells that expressed a control construct lacking the AH1 domain, demonstrating that membrane localization was required for pathway activity.

**Figure 2.**
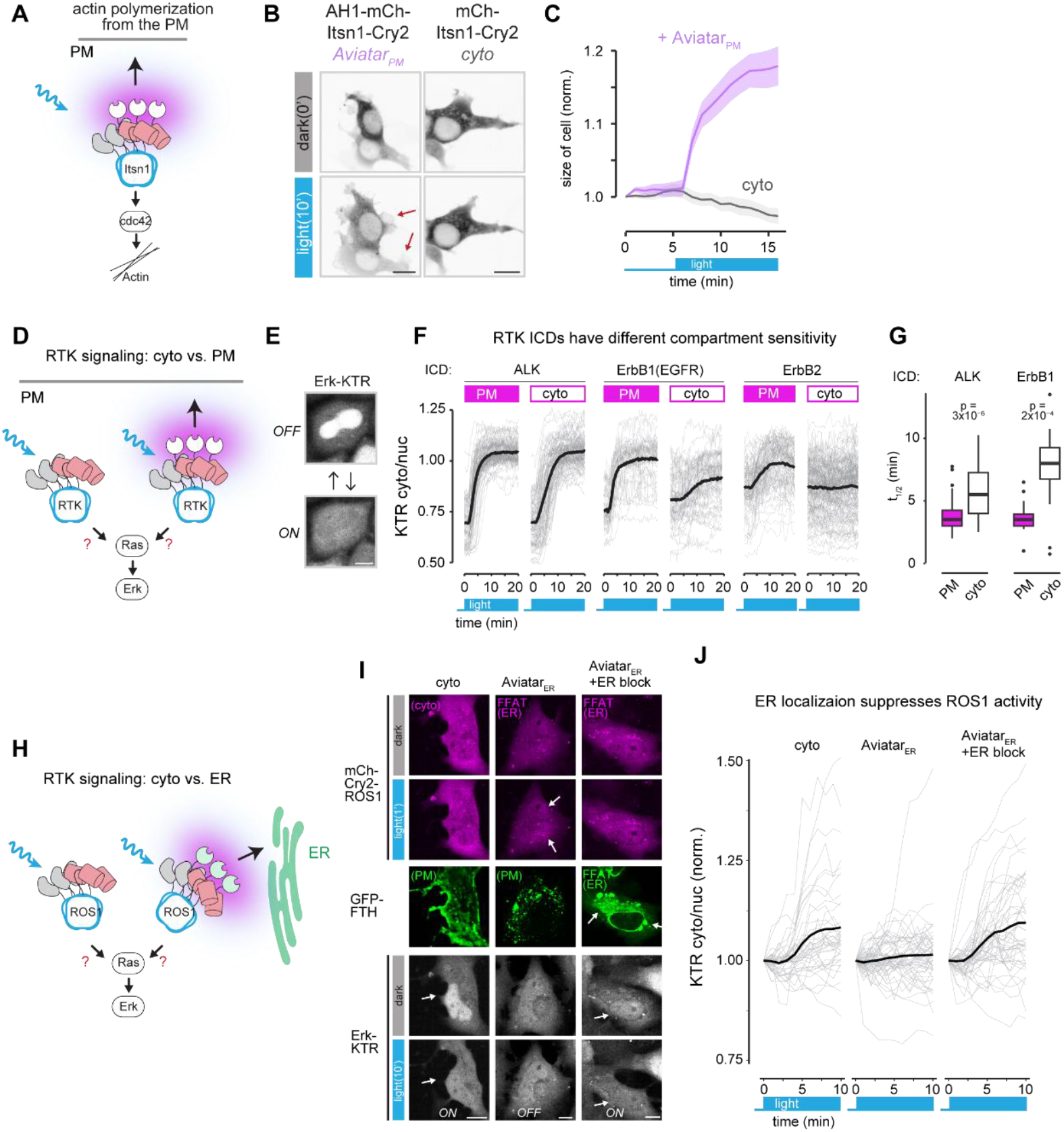
Aviatar can regulate signaling with specificity to subcellular compartments. **A**. Using Aviatar to control actin polymerization through recruitment of Cdc42-GEF ITSN1 DH-PH domain to the PM. **B**. Rapid filopodial and lamellipodial formation is observed when Itsn1 DH-PH domain is sent to PM compared to its clustering in the cytoplasm. **C**. Quantification of (**B**) showing mean +/-SEM of 32 cells (Aviatar) or 26 cells (cyto). Cell shape is normalized to time point 0 s. Blue light stimulation is performed after 5 minutes of imaging in dark. **D**. Comparing activation of receptor tyrosine kinase (RTK) intracellular domain (ICD) at either the PM or in cytosol upon clustering. All constructs include the Cry2olig variant. **E**. A kinase translocation reporter readout of Erk signaling (Erk-KTR) downstream of RTK signaling. The Erk-KTR reporter resides in the nucleus when Erk activity is off and translocates to the cytosol upon Erk activation. **F**. Quantification of Erk signaling (cytoplasmic/nuclear ratio of Erk-KTR). RTK ICDs were fused to either Stim1(PB)-Cry2olig (PM) or Cry2olig (cyto) and expressed in Hela cells. Grey traces represent single cells and black traces represent the mean. **G**. Half maximum activation time for ALK and ErbB1 ICDs in (**F**). Only cells with >0.1 KTR ratio change were quantified (40-80 cells per construct). **H**. Probing compartment specific effect on ROS1 ICD signaling in Beas2B cells. All constructs include the Cry2olig variant. **I**. Cry2olig-ROS1 or FFAT_Orp1_-Cry2olig-ROS1 were transfected to examine ROS1 signaling from the cytoplasm or ER, respectively. FFAT_Orp1_-FTH1 was co-transfected to block the Aviatar binding site on the ER, and PB_STIM1_-FTH1 was used as non-ER-targeting control for this blocking construct (PB_STIM1_ localizes to the PM). Arrows highlight the localization changes of either Aviatar_ER_-ROS1 or Erk-KTR after light stimulation. **J**. Quantification of Erk activity in (**I**). Grey traces represent tracked single cells, while black trace represents the mean. Scale bar, 10 µm.

### Aviatar reveals requirements for compartment localization for receptor tyrosine kinase fusions

We next use Aviatar to explore the consequences of differential localization of receptor tyrosine kinase (RTK) fusion activation. RTKs are normally a class of transmembrane proteins that, in response to stimulation by external ligands, multimerize and activate intracellular pathways that canonically originate at the membrane, including Ras/Erk and PI3K/Akt^37^. However, in certain cancers, chromosomal abnormalities generate RTK fusion oncoproteins: protein chimeras where the intracellular domain (ICD) of an RTK is fused to a multimeric region from an unrelated protein^38–40^. These fusions can reside in compartments other than the PM yet retain the ability to drive constitutive oncogenic signaling. Although > 20 distinct RTKs have been found as a part of fusions associated with cancer, certain RTKs appear as fusions more frequently than others. A recent study found the ICD of ALK in 53% of fusions. By contrast, the ErbB family, a well-studied family of 4 receptors (ErbB1-4) that is central in growth, development, and cancer ^41^, collectively was found in only ∼6%^38^. Additionally, ErbB family RTKs most often retain both their transmembrane domain and ICD in the context of the fusion, raising questions about whether these RTKs are more reliant on membrane localization compared to others.

We asked whether Aviatar could uncover compartment-specific differences in signaling among distinct RTK members (**Figure 2D**). We first examined the ALK ICD, comparing its activity when fused to Aviatar_PM_ (wBD: PB_STIM1_) vs. cytoplasmic clustering (no wBD) in HeLa cells. The stronger-clustering Cry2_olig_ was used here to ensure clustering of the large RTK payloads^17^. Signaling was assessed using a fluorescent biosensor of Erk activity (Erk-KTR) (**Figure 2E**)^42^. Surprisingly, both the PM-localized and cytoplasmic clustering of the ALK ICD signaled with comparable magnitude, with a slightly faster activation of Erk using Aviatar_PM_ (**Figure 2F-G**). We then performed the same experiment with the ErbB family of RTKs. In contrast to ALK, the ErbB1 (EGFR) ICD showed an increased magnitude of signaling from the PM relative to the cytoplasm, and PM-localized signaling was also faster to turn on (**Figure 2F, S3, S4**). On the other hand, ErbB2 could only be activated from the membrane but not from the cytoplasm, a pattern also observed for ErbB4. ErbB3 showed no activation either at the membrane or cytoplasm, consistent with ErbB3 being a pseudokinase ^12,43,44^. In sum, Aviatar reveals RTK-specific differences in the ability for activation from the cytoplasm vs the plasma membrane. These results also suggest that oncogenic ErbB family fusions often contain an intact transmembrane domain because, unlike ALK fusions, they require it to drive strong oncogenic signaling^38^.

We further validated Aviatar by localizing RTKs to an inhibitory — rather than stimulatory — compartment. Certain RTK fusions natively localize to compartments that suppress downstream signaling^45,46^. The CD74-ROS1 oncogene localizes to the ER and does not signal through the Ras/Erk pathway due to a suppressive environment at the ER, whereas cytoplasmic ROS1 fusions do stimulate Ras signaling^46,47^. We thus tested whether Aviatar_ER_ stimulation would suppress signaling from the ROS1 RTK (**Figure 2H**). ROS1 clustering in the cytoplasm triggered Erk signaling in lung epithelial Beas2B cells. By contrast, stimulation of Aviatar_ER_-ROS1 produced only minimal Erk activation, suggesting repression at the ER (**Figure 2I-J**). To confirm that signal suppression resulted from ER localization, we examined activation of Aviatar_ER_-ROS1 in cells where the ER-localized binding sites were pre-saturated with a decoy wBD, which we achieved with a constitutively multimerized FFAT_Orp1_ (GFP-FTH1-FFAT_Orp1_)^48^. In this condition, Aviatar_ER_-ROS1 did not visibly translocate to the ER in response to light, and Erk signaling was stimulated to levels comparable to cytoplasmic activation (**Figure 2I-J**).

Taken together, our results validate the Aviatar approach and demonstrate its ability to test consequences of signaling from varied cellular compartments.

### A universal GFP-targeting Aviatar for use with endogenously-labelled cells

A potential challenge of the Aviatar approach is identifying an appropriate wBD, one that is specific to the target compartment and is in the optimal range of affinity (minimal binding as a monomer, strong binding upon clustering). To address this challenge, we generated a universal Aviatar probe that would bind a common synthetic epitope, which could then be appended to a compartment specific protein for example through genomic tagging^49^. We thus generated an Aviatar against green fluorescent protein (GFP), which is commonly used to label endogenous proteins. To find an appropriate wBD, we tested a series of anti-GFP nanobodies with a range of binding affinities (K_d_ = 50, 310, 600, 3800 nM)^50^. When these candidate Aviatar_GFP_ probes were expressed in cells that harbored a membrane-anchored GFP (GFP-CAAX), all but the weakest-binding probe showed measurable membrane translocation upon light stimulation. Of the responsive probes, the weakest binder (LaG42, K_d_ = 600 nM) showed the lowest basal membrane binding in the dark while retaining strong translocation.

We then tested whether Aviatar_GFP_ could regulate translocation to stress granules (SGs), a compartment for which we could not identify a suitable wBD. SGs are dynamic biomolecular condensates of protein and RNA that form when cells are stressed^51,52^. We expressed Aviatar_GFP_ in HeLa cells where the stress granule scaffold G3BP1 was labelled with mClover3 (a derivative of GFP) at the endogenous locus, and we stimulated SG formation using heat stress^52^. SGs formed within 20 min of heating, indicated by large puncta of G3BP1. Although Aviatar_GFP_ remained largely diffuse in the dark, subsequent light stimulation drove colocalization of Aviatar_GFP_ with the SGs, with similar dynamic range for variants with binding affinities of 310 and 600 nM. Higher affinity Aviatar_GFP_ probes (K_d_ = 50 nM) similarly promoted recruitment to SGs but also showed elevated basal association to SGs (**Figure S5**). Thus, Aviatar_GFP_ provides a generalizable implementation of the strategy that can be applied to numerous established cell lines and protein fusions, with a range of affinities to optimize for specific applications.

## Discussion

Aviatar is a general and straightforward strategy to drive the localization of individual proteins towards user-defined cellular compartments by regulating binding avidity. Although avidity regulation is commonly observed in natural systems^15,21,23–25,29,31,53^ and plays a role in a handful of prior optogenetic probes ^11,12,20,54,55^, here we find that this principle can be explicitly harnessed for the systematic generation of a variety of translocation probes.

Aviatar’s advantage over existing approaches is its simplicity. As a single component switch, only one protein must be optimized. This obviates the need for tuning relative expression levels between multiple system components, which is impractical in most experimental settings beyond single cells. Single-component systems also simplify delivery because the entire system can be encoded within one short sequence of RNA or DNA. Binding endogenous targets minimizes the risk of inadvertently altering the function of the target compartment by expressing constitutive anchors. Simplicity also derives from its modularity -- target compartments or effector domains can be exchanged without disruption of the inducer module, allowing rapid testing and optimization of these components. This contrasts with alternative methods which, while powerful, rely on domain insertion of an allosteric switch like *As*LOV2^9,56–58^, a strategy that is highly sensitive to insertion position and thus requires laborious screening^59^.

Importantly, Aviatar probes have two inextricable outputs: translocation and protein clustering. For many applications, this offers a compact encoding of two useful functions, for example for stimulating RTK signaling^12^. For other targets like ITSN1/Cdc42, clustering is not necessary but also not detrimental for activation^3,34^. In certain cases, however, clustering can suppress activity^60,61^. The essentiality of clustering must also be considered when the *target* is clustered, which in principle could trigger Aviatar translocation even in the absence of the inducer. Nevertheless this concern can be minimized with selection of an appropriately weak wBD, as we demonstrate with minimal basal binding of AviatarGFP to stress granules using weak *α*GFP nanobodies.

Aviatar is a general strategy for inducible translocation. While we exclusively used optogenetic clustering, Aviatar’s modularity allows easy expansion of its inputs, for example to chemicals, temperatures, or other platforms for optogenetic clustering^13,52,62–65^. The continued progress in engineered protein and peptide binders, as well as in GFP-labelling of endogenous proteins, will allow Aviatar to navigate to an expanding set of applications.

## Acknowledgements

This work was supported by funding from the National Institutes of Health (R35GM138211 for L.J.B.), the National Science Foundation (CAREER 2145699 to L.J.B.), and the Penn Center for Precision Engineering for Health (CPE4H Pilot grant to L.J.B.). Cell sorting was performed on a BD FACSAria Fusion that was obtained through NIH S10 1S10OD026986.

## Contributions

Z.D.H. and L.J.B. conceived the study. Z.D.H., Y.G, and A.B. tested all Aviatar probes. Z.D.H., Y.G. applied Aviatar for controlling cytoskeleton. Z.D.H., Y.G., Y.C.G., H.B. characterized Aviatar-RTK subcellular signaling. Z.D.H., Y.G., Y.C.G., analyzed the imaging data. L.J.B. supervised the work. Z.H. and L.J.B. wrote the manuscript and made figures, with editing from all authors.

## Methods

### Molecular cloning

All plasmids were constructed in the pHR backbone with a CMV promoter. Plasmids were assembled using restriction endonuclease digestion and PCR with Q5 polymerase (NEB M0493S), followed by HiFi assembly (NEB E2621S). The resulting reaction was transformed into chemically competent cells (NEB Turbo, C2984H). Cry2, Alk, ErbB1-4 were sourced from previous work ^12,13,66^. Ros1 was a gift from Dr. Nidhi Sahni (MD Anderson). Weak binding domains were synthesized from IDT, and GFP nanobodies^50^ were a gift from Dr. Michael Rout (Rockefeller University). All constructs containing Cry2 were fused to mCherry.

### Cell culture and cell lines

HEK 293T and HeLa cells were cultured in DMEM with 10% fetal bovine serum (FBS) (Mediatech 35-010-CV)and 1% penicillin/streptomycin (P/S). Beas2B cells were cultured in RPMI with 10% fetal bovine serum (FBS) and 1% penicillin/streptomycin (P/S). All cells were incubated at 37 °C with 5% CO_2_ during imaging. Before testing RTK signaling, cells were starved overnight in media without FBS.

RTK signaling was tested in HeLa and Beas2B cell lines stably expressing Erk-KTR-iRFP as described previously (**Figure 2D-I**)^66^. HEK 293T cells stably expressing iRFP-Kras(CAAX) and GFP-Kras(CAAX) were described previoysly^13,36^. HeLa cells with endogenously labeled G3BP1-mClover3 were a gift from Dr. Ophir Shalem (Children’s Hospital of Philadelphia).

### Protein expression

All the Aviatar components were transiently transfected into cells using Fugene4K (Promega, E5911). Cells were seeded one day before transfection. 6-12 hours after transfection, cells were washed with fresh media.

### Confocal imaging

Live-cell imaging was performed using a Nikon Ti2-E spinning disk (Yokagawa CSU-W1) confocal microscope equipped with 405/488/561/640nm laser lines and an sCMOS camera (Photometrics). CO_2_ and ambient temperature were under control of an environmental chamber (Okolabs). Cells were imaged with either a 20X, 40X air, or 60x oil objective. Aviatar constructs were stimulated using the 488 nm laser.

All imaging experiments were performed under standard cell culture conditions (37 °C, 5%CO_2_) except for data in **Figure 3D-F**. These experiments were performed with the thermoPlate, a device that precisely and rapidly controls the temperature in 96-well plates^52^.

**Figure 3.**
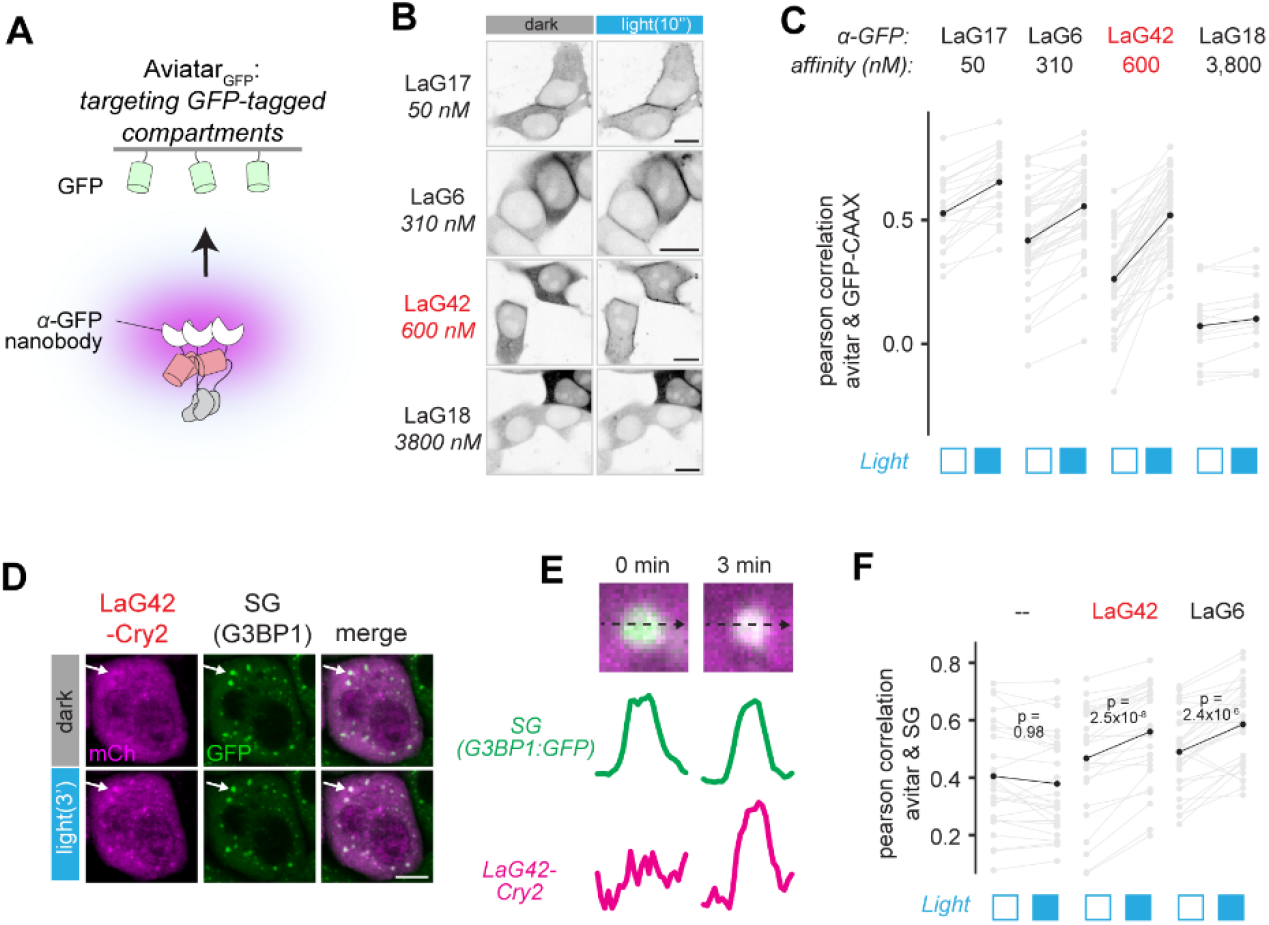
Targeting GFP-tagged compartments with Aviatar_GFP_. **A**. Schematic diagram of the Aviatar_GFP_. **B**. Screening α-GFP nanobodies for suitability. HEK 293T cells stably expressing GFP-CAAX_KRAS_ were transfected with Cry2 fused to α-GFP nanobodies (LaG), with K_d_ as indicated. **C**. Quantification of Aviatar_GFP_ translocation to membrane-bound GFP in (**B**) using Pearson correlation between GFP and Aviatar_GFP_ intensity. Grey traces represent tracked single cells. Black traces show mean of grey traces. **D**. Selected Aviatar_GFP_ probes are transfected in HeLa cells with G3BP1 endogenously labelled with mClover3, a GFP derivative. Cells were preincubated at 43 °C (heat stress) to induce the formation of stress granules (SG), following which cells were stimulated with blue light. Arrows highlight an SG with increased Aviatar_GFP_ localization after light stimulation. **E**. Intensity profile of the highlighted SG in (**D**). **F**. Pearson correlation of Aviatar_GFP_ and SG intensity is used to quantify colocalization. Scale bar, 10 µm.

Before imaging Erk-KTR, cells were stained with 5 µg/mL Hoechst 33342 Hydrochloride (Cayman 15547) for 15 min followed by fresh media washes.

### Image analysis

#### a. Membrane recruitment

The plasma membrane association of Aviatar in **Figure 1B-I** was calculated using normalized plasma membrane (PM)/cytoplasmic mean fluorescence ratio. Cells with iRFP-Kras(CAAX) marker were segmented using the MorpholibJ module in ImageJ^67^. These segmented images were then imported into a CellProfiler script, in which the cells with positive mCherry signal were selected. The PM is defined by a 3 pixel ring inside the edge of the contour of each cell. The PM and whole cell fluorescence were then quantified. The degree of membrane recruitment is quantified using the mean intensity of PM divided by the mean intensity of the whole cell.

#### b. Colocalization

Cells with expression of both Aviatar and an organelle marker were segmented manually using a customized MATLAB script and the intensity of each pixel in each channel was extracted^8^. The colocalization between 2 different fluorescent signals in this manuscript is quantified using Pearson correlation quantification in R. The intensity profiles were exported from ImageJ.

#### c. Erk-KTR translocation

Cells were segmented with nuclear stain (Hoescht) using CellProfiler, then tracked using nucleus location using a customized script in R. Only tracks starting from 0 and lasting at least 15 minutes were kept. The mean intensity of the cytoplasm was estimated by a 3 pixel ring surroundng the nucleus. The cytoplasm/nucleus ratio was then calculated for visualization.

## Figures and legends

**Figure S1.**
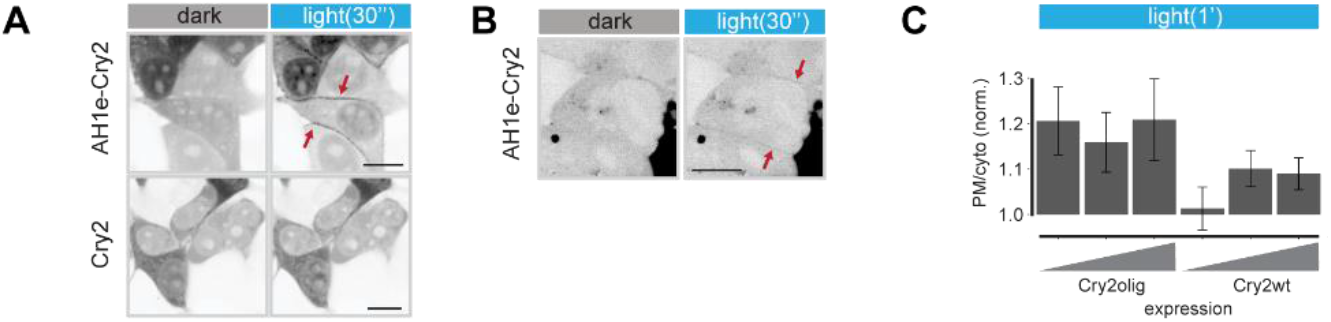
Aviatar_PM_ works at even low expression levels in HEK 293T cells. **A**. Representative images showing light stimulation of Aviatar under low expression levels where clusters of Cry2 alone cannot be observed. **B**. Representative images showing light stimulation of cells with low expression levels that are minimally above background. **C**. Comparison of Aviatar_PM_ binding capacity when using Cry2wt vs Cry2olig, which has a stronger propensity to cluster. Scale bar, 10 µm. All Aviatar_PM_ constructs in this figure use AH1 as the wBD.

**Figure S2.**
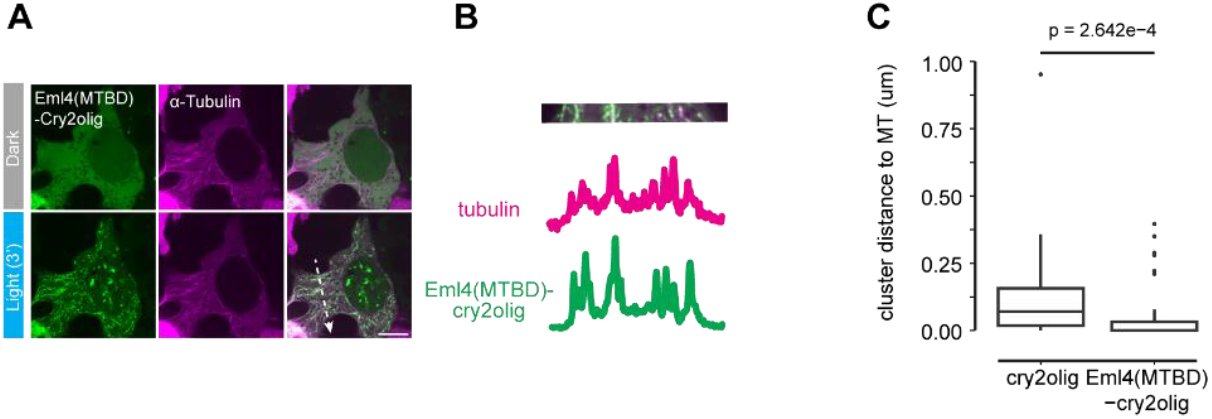
Microtubule targeting with Aviatar using truncated EML4 as the wBD fused with Cry2olig. **A**. Representative images showing light stimulation of Aviatar_MT_ in HEK 293T cells. α-Tubulin is used as a microtubule marker. **B**. Intensity profile of area highlighted with dashed arrow in (**A**). **C**. Quantification of the 3 min time point of (**A**). Scale bar, 10 µm.

**Figure S3.**
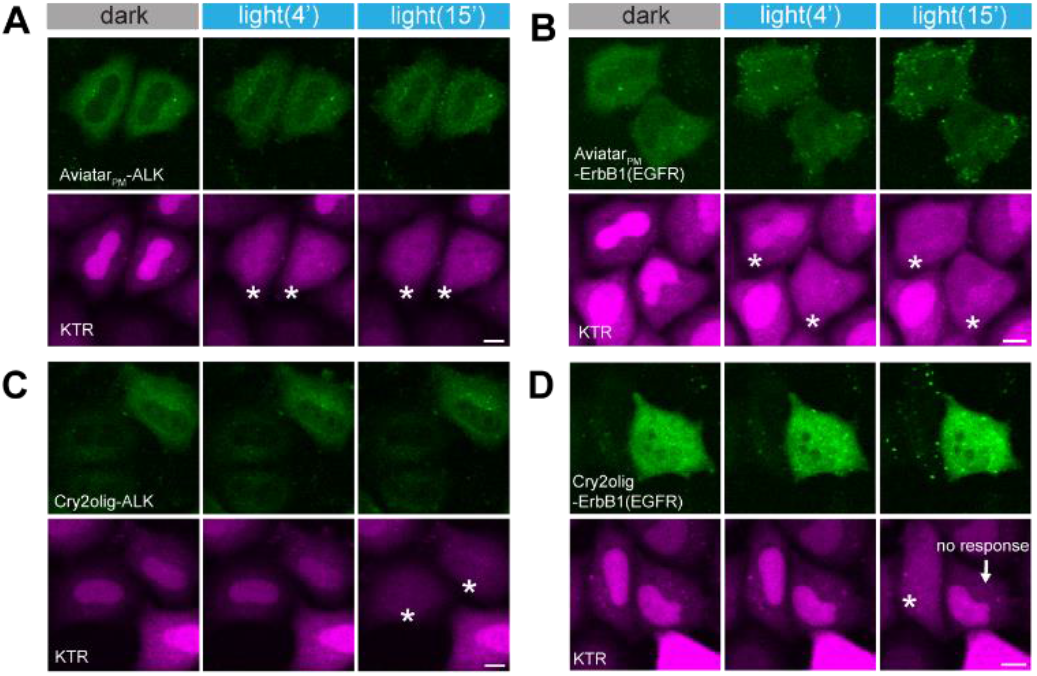
Representative images of Figure 2F. **A**. Representative images showing light stimulation of Aviatar_PM_-ALK in Hela cells stably expressing Erk-KTR. **B**. Representative images showing light stimulation of Aviatar_PM_-ErbB1(EGFR) in Hela cells stably expressing Erk-KTR. **C**. Representative images showing light stimulation of Cry2olig-ALK in Hela cells stably expressing Erk-KTR. **D**. Representative images showing light stimulation of Cry2olig-ErbB1(EGFR) in Hela cells stably expressing Erk-KTR. Cells with KTR translocation are labeled with stars. Scale bar, 10 µm.

**Figure S4.**
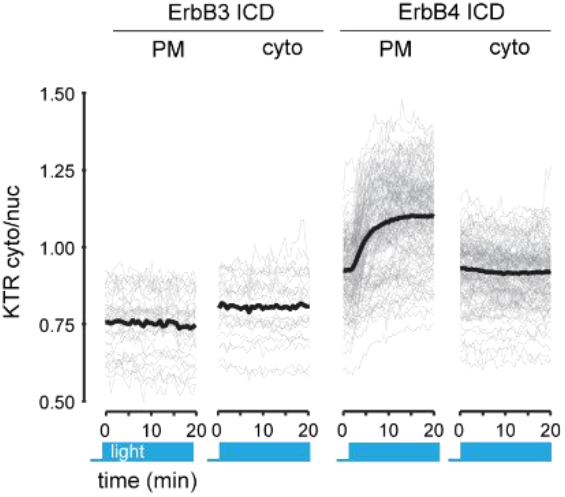
Cytoplasmic vs PM activation of ErbB3 and ErbB4. Quantification as described in Figure 2F.

**Figure S5.**
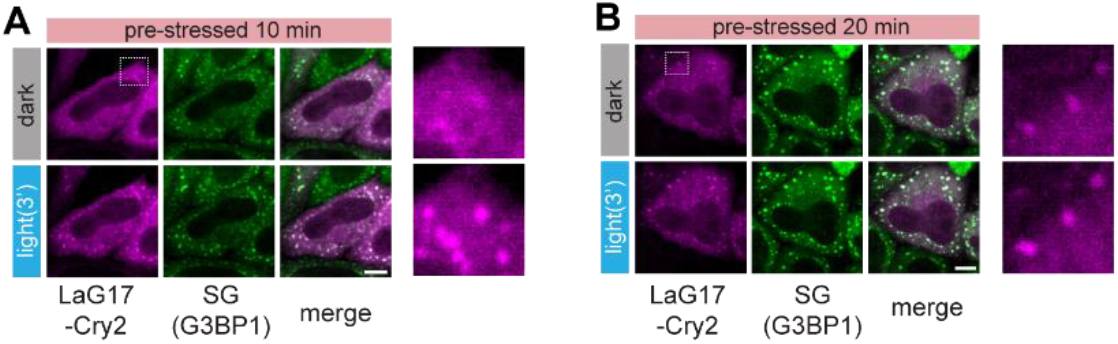
Aviatar_GFP_ displays different dynamic range according to avidity of GFP. Aviatar_GFP_ (LaG17) was transfected in Hela cells with endogenously tagged G3BP1-mClover3 (a GFP variant). Cells were stimulated with blue light after different time of incubation at 43°C. G3BP1 enriched stress granules (SG) are larger and brighter with longer incubation time. The regions in dashed box were zoomed in on the right. **A**. Cells were pre-stressed for 10 minutes before light stimulation. **B**. Cells were pre-stressed for 20 minutes before light stimulation. Scale bar, 10 um.

**Movie S1. Light-induced Aviatar**_**PM**_ **translocation to the plasma membrane, using PB**_**STIM1**_ **as the wBD**. Time, mm:ss. Blue box indicates light stimulation. Scale bar, 10 um.

**Movie S2. Light-induced Aviatar**_**PM**_ **translocation to the plasma membrane, using PH**_**Dnm1**_ **as the wBD**.Time, mm:ss. Blue box indicates light stimulation. Scale bar, 10 um.

**Movie S3. Light-induced Aviatar**_**Endosome**_ **translocation to endosomes, using FYVE**_**Fyco1**_ **as the wBD**.Time, mm:ss. Blue box indicates light stimulation. Scale bar, 10 um.

**Movie S4. Light-induced Aviatar**_**Golgi**_ **translocation to endosomes, using PH**_**Fapp1(R18L)**_ **as the wBD**. Time, mm:ss. Blue box indicates light stimulation. Scale bar, 10 um.

**Movie S5. Light-induced Aviatar**_**ER**_ **translocation to endosomes, using FFAT**_**Orp1**_ **as the wBD**. Time, mm:ss. Blue box indicates light stimulation. Scale bar, 10 um.

**Movie S6. Light-induced Aviatar**_**MT**_ **translocation to microtubules, using KIF5B(motor)**_**1**_ **as the wBD**. Time, mm:ss. Blue box indicates light stimulation. Scale bar, 10 um.

**Movie S7. Optogenetic stimulation of actin polymerization using Aviatar**_**PM**_**-DHPH**_**Itsn1**_ **translocation**. Time, mm:ss. Blue box indicates light stimulation. Scale bar, 10 um.

**Movie S8. Aviatar**_**GFP**_ **translocation to the PM in cells stably expressing GFP-Kras(CAAX), using LaG42 as the wBD**. Time, mm:ss. Blue box indicates light stimulation. Scale bar, 10 um.

**Movie S9. Aviatar**_**GFP**_ **translocation to stress granules in cells where G3BP1 is tagged with mClover3, using LaG42 as the weak binding domain**. Time, mm:ss. Blue box indicates light stimulation. Scale bar, 10 um.

